# Methylation of RNA Cap in SARS-CoV-2 captured by serial crystallography

**DOI:** 10.1101/2020.08.14.251421

**Authors:** M. Wilamowski, D.A. Sherrell, G. Minasov, Y. Kim, L. Shuvalova, A. Lavens, R. Chard, N. Maltseva, R. Jedrzejczak, M. Rosas-Lemus, N. Saint, I.T. Foster, K. Michalska, K.J.F. Satchell, A Joachimiak

## Abstract

The genome of the SARS-CoV-2 coronavirus contains 29 proteins, of which 15 are nonstructural. Nsp10 and Nsp16 form a complex responsible for the capping of mRNA at the 5′ terminus. In the methylation reaction the S-adenosyl-L-methionine serves as the donor of the methyl group that is transferred to Cap-0 at the first transcribed nucleotide to create Cap-1. The presence of Cap-1 makes viral RNAs mimic the host transcripts and prevents their degradation. To investigate the 2′-O methyltransferase activity of SARS-CoV-2 Nsp10/16, we applied fixed-target serial synchrotron crystallography (SSX) which allows for physiological temperature data collection from thousands of crystals, significantly reducing the x-ray dose while maintaining a biologically relevant temperature. We determined crystal structures of Nsp10/16 that revealed the states before and after the methylation reaction, for the first time illustrating coronavirus Nsp10/16 complexes with the ^m7^GpppA_m2′-O_ Cap-1, where 2′OH of ribose is methylated. We compare these structures with structures of Nsp10/16 at 297 K and 100 K collected from a single crystal. This data provide important mechanistic insight and can be used to design small molecules that inhibit viral RNA maturation making SARS-CoV-2 sensitive to host innate response.

## INTRODUCTION

Five coronaviruses can induce clusters of severe respiratory diseases in humans: 229E, OC43, SARS, MERS and SARS-CoV-2 ^1,2^. The outbreak of Severe Acute Respiratory Syndrome (SARS-CoV) in 2003 and Middle East Respiratory Syndrome (MERS-CoV) in 2012, showed a high fatality rate of 11% and 34%, respectively, but had limited geographical spread ^3,4^. In contrast, in the past 7 months SARS-CoV-2 (the cause of COVID-19) has spread rapidly around the world and infected millions of people, killing over half a million. Though, it has a significantly lower fatality rate (estimated at 0.5 – 2%) than MERS and SARS the amount of people infected is hundreds of times higher, and it caused significant economic hardship and extraordinary social restrictions. In many countries, after six months from the first reports of COVID-19, the number of cases is still increasing ^5^. In the absence of herd immunity, vaccines, or drugs against the virus, the situation is frightful. Since the beginning of the worldwide outbreak of COVID-19, the scientific community has joined efforts to understand SARS-CoV-2 biology and to find chemical compounds that either block virus replication or enhance human immunological response. Initial focus on repurposing existing drugs that potentially could provide immediate treatment has thus far resulted only in limited success. Finding inhibitors of a biological cycle of SARS-CoV-2 or vaccines is, therefore, critical for global health, safety, and wellbeing.

SARS-CoV-2 β-coronavirus has a large (~ 30 kb) and complex (29 proteins) (+) sense single-stranded RNA genome ^6^. The RNA present in the mature virion resembles human mRNA: (i) it is capped on its 5′-end, (ii) contains a 3′-poly-A tail, and (iii) after infection can be directly translated to the two polyproteins Pp1a and Pp1ab using host machinery. These polyproteins are then matured into 15 non-structural proteins (Nsp), that assemble into a large replication-transcription complex. The RNA is also used as a template for biosynthesis of (-) sense RNA that serves to make additional copies of (+) sense RNA and several sub-genomic RNAs for the translation of four structural and 9-10 accessory proteins ^6^. For these RNAs to serve as mRNAs, they must undergo post-transcription modifications to resemble human mRNA. The RNA maturation involves several enzymatic steps performed by viral Nsps; Nsp13 is a bifunctional RNA/NTP triphosphatase (TPase) and helicase; Nsp14 is a bifunctional 3′-5′ exonuclease and guanine N7 methyltransferase; and Nsp16 is Mg^2+^ dependent ribose 2′-O methyltransferase and an elusive guanylyltransferase.

In eukaryotes, the attachment of the ^m7^G Cap at the 5′-end of RNA protects the transcript from mRNA turnover in the pathway dependent on 5′ → 3′ exoribonucleases activity, such as XRNs ^7^, and is required for several cellular processes: maturation of mRNA, pre-mRNA nuclear export, and protein synthesis ^8^. The Cap at the 5′ terminus of eukaryotic transcripts starts with 7-methylguanosine at the reversed position and is connected through an uncommon 5′ to 5′ triphosphate bridge, which is attached to a nucleotide that undergoes methylation at 2′-O-ribose. The canonical pathway of the mRNA capping mechanism requires the subsequent action of multiple enzymes ^8^. At first, RNA 5′-triphosphatase (RTPase) cleaves the 5′-terminal γ-β phosphoanhydride bond of the nascent mRNA and enables further modification of the newly synthesized transcript at its 5′-terminus. Secondly, the diphosphate mRNA is capped using GMP by guanylyltransferase (GTase). Next, the (guanine-N7)-methyltransferase adds the methyl group to the 5′-terminal guanosine creating a precursor: Cap-0. Additionally, the 2′-O methyltransferase transfers the methyl group to the 2′-O-ribose of the first transcribed nucleotide at mRNA, making the ^m7^GpppN_m2′-O_ (Cap-1) structure ^9^. S-adenosyl-L-methionine (SAM) is a donor of the methyl group and is converted to S-adenosyl-L-homocysteine (SAH) during the reaction ^10^. In humans and other higher eukaryotes, the second nucleotide of mRNA then undergoes methylation to produce Cap-2 ^11^.

The post-transcriptional modification of RNA is also essential for efficient translation of viral transcript in a eukaryotic host ^12^. Influenza, Ebola, measles, poxvirus, and coronaviruses attach Cap to their genomic and sub-genomic RNA transcripts to mimic host molecules and escape from the innate immunological response ^13^. Studies done in the last decade show that the 2′-O methylation of the Cap-0 facilitates this process by preventing activation of type I interferon that is induced by the cytoplasmic RNA sensors, melanoma differentiation-associated protein 5 (MDA5), and retinoic acid inducible gene-I (RIG-1) ^14–16^. Mutations in 2′-O methyltransferase render the viruses sensitive to the interferon-inducible immunological pathways ^17^.

Methyltransferases (MTases) are common in all metazoan and viral genomes ^18^. SARS-CoV-2 possesses two MTases, Nsp14 and Nsp16, both of which form a heterodimer complex with Nsp10 ^19,20^, which is a critical protein for virus replication and fidelity ^21^. The sequence similarity of viral 2′-O MTases is high, suggesting common origin. Sequence identity of Nsp16 between SARS-CoV-2 and SARS-CoV-1 is 95%, while the identity of SARS-CoV-2 Nsp16 to the MERS-CoV enzyme is 66% ^20^. The active site of Nsp16 MTase is conserved in the *Coronaviridae* family; they all utilize a Lys-Asp-Lys-Glu catalytic tetrad essential for the enzymatic activity ^22,23^.

It has been shown that the Nsp10/16 complex methylates the Cap-0 (^m7^GpppA_2′-OH_) to form Cap-1 (^m7^GpppA_m2′-O_) by adding a methyl group to the ribose 2′-O of the first nucleotide (usually adenosine, in CoVs) of the nascent mRNA using SAM as the methyl group donor ^24^. The Nsp16 activity is necessary for Nsps’ translation from new copies of viral (+) sense RNA as well as structural and accessory proteins translation from sub-genomic RNA which are all transcribed from the viral (-) sense RNA template. Vaccination with Nsp16 defective SARS-CoV or an immunogenic disruption of the Nsp10/16 interface protects mice from a lethal SARS challenge ^25^. Therefore, blocking Nsp16 activity should reduce viral proliferation, making the protein an attractive drug target.

While several structures of Nsp10/16 in a complex with Cap-0 have been determined by crystallography (PDB entries: 6WQ3, 6WRZ, 6WVN, 6WKS), none have captured the post-methylation state. We conducted serial synchrotron crystallography (SSX) experiments at 297 K to test whether low radiation dose could help uncover the structure of Nsp10/16 in a complex with Cap-1. Previously, x-ray free-electron laser (XFEL) serial crystallography (SFX) had established methods to study molecular dynamics and chemical reactions in protein crystals occurring at femtosecond to millisecond timescales ^26–28^. The XFEL-based protocol enabled to analyze protein structures with no radiation damage, in the so-called “diffraction before destruction” mode ^29,30^. SFX has been successfully applied to determine the structure of the membrane protein complex of photosystem I and of several other macromolecules using time-resolved approach ^29^. The synchrotron equivalent, SSX, uses a similar approach to SFX but can access longer, biologically relevant time scales. Moreover SSX is a much more accessible technique at a number of light sources and less sample consuming than SFX ^31^. Nonetheless, delivering hundreds of thousands of batch-grown crystals to the x-ray beam still presents challenges. Two approaches are common: the first uses a liquid injector of microcrystals ^29,32,33^, and the second uses a fixed-target system where crystals are deposited on a chip and scanned through the x-ray beam ^34,35^. Here we present three crystal structures of the Nsp10/16 complex determined by fixed-target SSX at 297 K: in the presence of SAM, Cap-0/SAM, and with Cap-1/SAH generated by metal-dependent conversion of Cap-0/SAM substrates. We compare these structures with structures of Nsp10/16 at 297 K and 100 K collected from a single crystal. We observe the state of the molecules in the crystal before and after methylation. The uniqueness of the Cap-1 structure shows the advantages of the SSX method in structural studies: allowing the use of larger crystals, reducing radiation damage, and being closer to physiological temperatures.

## RESULTS AND DISCUSSION

### Serial crystallography for the Nsp10/16 2′-O MTase from SARS-CoV-2

We collected SSX data for the Nsp10/16 crystals using the fixed-target SSX system (Advanced Lightweight Encapsulation Crystallography (*ALEX)* mesh-holder) as depicted in Figure 1A and Extended Data Fig. 1. A crystal slurry is deposited on a nylon mesh, which immobilizes the crystals; they are then encapsulated between two polyester films ^36^. A representative diffraction image recorded from a single Nsp10/16 crystal illustrates the quality of the obtained data (Extended Data Fig. 2B). We collected data from seven chips with ~ 36,100 images per chip with an exposure time of 50 ms at 100% x-ray transmission. Including overheads, motor acceleration, deceleration, and settling time, each mesh was in the beam for approximately 90 minutes, collecting images at a rate of 7 Hz. SSX experiments were conducted on three separate days: first to collect data on Nsp10/16/SAM complex. Second for Nsp10/16/SAM crystals with SAM that were soaked with Cap-0 in the presence of EDTA. Third to collect data for the Nsp10/16/Cap-1/SAH complex, resulting from the *in crystallo* reaction converting Cap-0/SAM to Cap-1/SAH (Fig. 1B). As discussed below, this process occurred in the crystals in the presence of 1 mM Mg^2+^ and the reaction is significantly slowed down by addition of EDTA.

**Figure 1.**
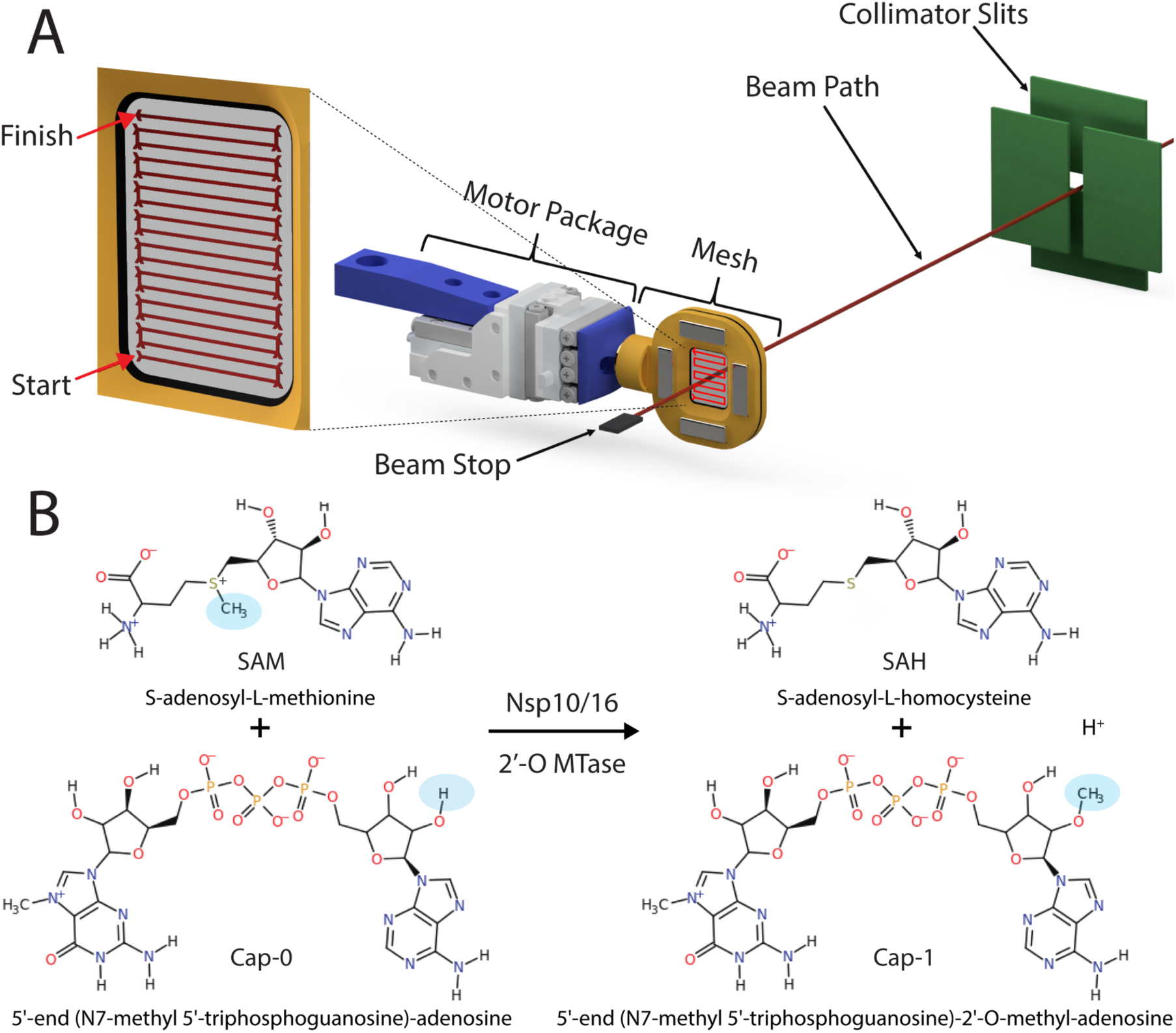
Fixed-target serial crystallography data collection on Nsp10/16 crystals from SARS-CoV-2. System for SSX with moving crystal holder developed at APS beamline 19-ID (**A**). Chemical formula of reaction done by Nsp10/16 2′-O methyltransferase, transfer of methyl group labeled as blue oval (**B**).

For initial data processing, we used the *Kanzus* automated data pipeline. *Kanzus* integrates the 19-ID data collection system with the Argonne Leadership Computing Facility (ALCF), using the Theta supercomputer for high-speed on-demand data analysis. Extended Figure 3A presents an image with labeled indexed reflections and shows that most observed reflections of Nsp10/16 crystals were indexed. PRIME identified inadequate low-resolution statistics from the integration files, thus we subjected them to outlier rejection. Integration files with resolution lower than 3.5 Å, containing less than 90 reflections, or with a unit cell volume 4.5% different from the average were removed from the scaling process (Extended Data Tab. 1, Fig. 2C, 2D). The total number of integrated files that went into PRIME for Nsp10/16 bound with SAM, Cap-1/SAH, and Cap-0/SAM were 7,951; 5,038 and 4,704 respectively. The overall hit-rate (images used for structure divided by images taken) for the five meshes was 7%. Some positions on the mesh had multiple crystals, 11% of ‘hits’ had two lattices, and fewer than 1% had three (Extended Data Tab. 1, Fig. 3B). For structure determination we used all indexed lattices in each diffraction image. Two structures from SSX were solved at 2.25 Å (Nsp10/16/SAM and Nsp10/16/Cap-1/SAH), and a third at 2.18 Å (Nsp10/16/Cap-0/SAM) (Extended Data Tab. 1). The highest resolution shell in PRIME analysis was specified using a CC_1/2_ cut-off below 0.40. Completeness for the SSX data was close to 100% and R_work_ was 20.65%, 22.31%, and 21.70% for Nsp10/16/SAM, Nsp10/16/Cap-1/SAH, and Nsp10/16/Cap-0/SAM, respectively (Extended Data Tab. 2). For a comparison to SSX data, we also collected data from a single crystal of the Nsp10/16 with Cap-0/SAM and Cap-1/SAH in a capillary at 297 K. These data could only beprocessed to 2.65 Å with reasonable statistics (Extended Data Tab. 1, 2). Using RADDOSE-3D ^37^, we estimated the accumulated dose of x-ray for Nsp10/16 from SSX to be 0.12 MGy, whereas the dose for the capillary-mounted crystal was 4.97 MGy over 40 times higher.

### 297 K structure of the Nsp10/16 heterodimer from SARS-CoV-2

The Nsp10/16 complex is an α/β heterodimer (Fig. 2). The 139 amino acid long Nsp10 has unique fold formed by 5 α-helices and a pair of antiparallel β-strands which are facing Nsp16 in the complex. Nsp10 possesses two zinc ions coordinated by the CCHC and CCCC motifs that are 100% conserved in β-coronaviruses ^20^. Nsp10 is involved in forming the complexes with multiple SARS-CoV-1 and −2 proteins. The best characterized is the complex with Nsp14 3′-5′ exoribonuclease (and guanine N7 methyltransferase) and Nsp16 that is 2′-O-ribose methylase. Previous research showed that Nsp16 is only active in the presence of Nsp10 ^19^. Interestingly, the activity of 2′-O MTase of the Nsp10/16 complex was reduced by using a peptide compound based on the Nsp10 sequence ^38^. Nsp10 is a molecular co-factor for various SARS-CoV-2 enzymes and therefore a good candidate for design of molecules that will affect its structure or disrupt the interaction with Nsp14 or Nsp16 2′-O MTases.

**Figure 2.**
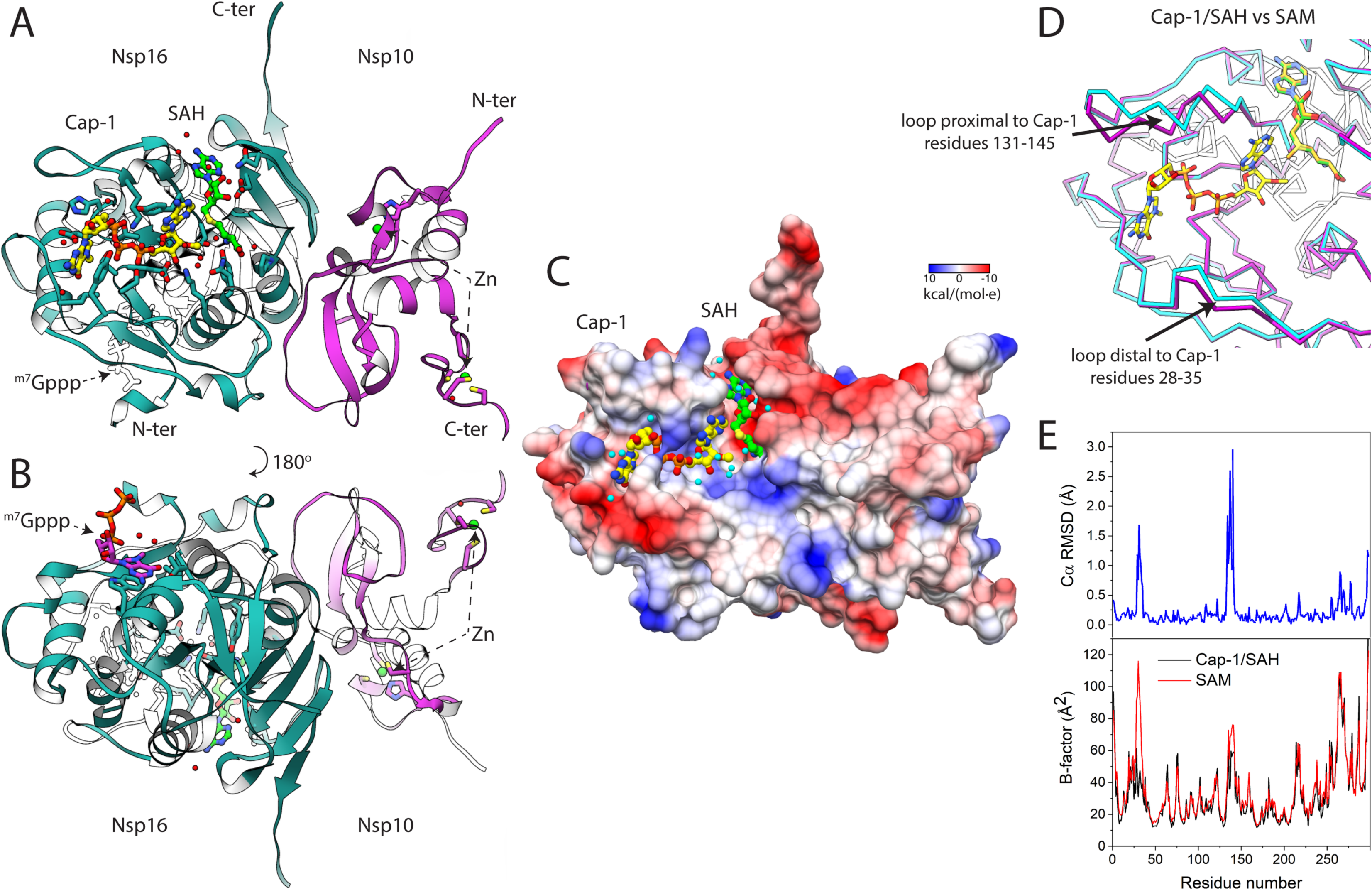
297 K crystal structure of the Nsp10/16 in a complex with ^m7^GpppA_m2′-O_ (Cap-1) and SAH determined using fixed-target SSX. The heterodimer assembly of the Nsp10/16 complex (**A, B**); Nsp10 (pink), Nsp16 (blue). The 2′-O MTase active site with Cap-1 and SAH (**A**). The potential allosteric site of Nsp16 with bound ^m7^Gppp (**B**). The Coulombic electrostatic potential on the surface of the Nsp10/16 heterodimer in complex with Cap-1/SAH calculated at 298 K using default settings in UCSF Chimera (**C**). Comparison of 297 K crystal structures of the Nsp10/16 complex with Cap-1 and SAH (blue – PDB entry 7JHE) superimposed to the structure with SAM (pink – PDB entry 6XKM) (**D**). Binding of mRNA Cap to Nsp16, plots of RMSD (top) and B factor between structures with SAM and Cap-1/SAH (**E**).

The structure of Nsp16 is characterized by a Rossmann fold, with a large β-sheet surrounded by α-helices, β-strands and loops. Nsp16 contains 11 α-helices, 12 β-strands and 7 loops. Nsp16 has a centrally positioned β-sheet (β3↑,β2↑,β1↑,β4↑,β5↑,β7↓,β6↑) with only one antiparallel strand β7. The SAM and the Cap binding sites are located at the surface of Nsp16. None of the Nsp10 residues have a direct connection with the ligands’ binding sites (Fig. 2A, 2C), but Nsp10 is required for Nsp16 MTase activity ^21^. Additionally, we observed partial electron density for the 7-methyl-guanosite-5′-triphosphate (^m7^Gppp) of the Cap in the possible allosteric site of the Nsp10/16 complex (Fig. 2B). Analysis of the surface electrostatic charges of the Nsp10/16 structure shows positively charged residues that form an elongated binding pocket for Cap, with approximate length of 21 Å (Fig. 2C). In proximity of the Cap-0 methylation site there is an elongated positively charged surface that can potentially bind longer RNA (Fig. 2C). Moreover, there are reports that two zinc fingers of Nsp10 contribute to non-specific binding of nucleic acids ^24^, which could further indicate that Nsp10/16 not only binds the Cap, but also interacts with longer RNA fragments.

The SSX experiments on the Nsp10/16 crystals revealed three states: one with SAM bound, second with SAM and Cap-0 in the active site prior reaction occurring, and the third state after methyl transfer with Cap-1 and SAH. The electron density clearly showed the methyl group present on the adenosine moiety at the 2′-O-ribose position of the Cap-0 and no methyl present on SAH (Fig. 4C). Moreover, the 297 K single crystal structure has also clear electron density for the methylation site of the Cap-0 2′-O-ribose, but with partial occupancy.

The structures determined at 100 K using single crystal and at 297 K using SSX obtained in the same crystal forms are very similar, as illustrated by superpositions of our Nsp10/16/Cap-1/SAH structure with the complex of Cap-0/SAH (PDB entry 6WQ3) (Extended Data Fig. 4). For these structures, the overall root mean square deviation (RMSD) of Cα atoms is 0.43 Å. We observed less than an 1.1 Å shift of the loops that form the binding sites for the Cap and zinc ions. Comparison of the SSX structures of Nsp10/16 with Cap-0/SAM and Cap-1/SAH does not reveal any significant changes in protein backbone (Cα RMSD: 0.52 Å) (Extended Data Fig. 4B). On the other hand, the comparison of Nsp10/16/SAM and Nsp10/16/Cap-1/SAH complexes determined using SSX indicate larger conformation changes in the Nsp16 that are induced by the Cap-0 binding and Cap-1 formation. We observed movement of two loops (comprising 28-35 and 131-145 residues of Nsp16) which form the mRNA Cap binding groove; positions of some residues are shifted as much as 3.0 Å, with an overall Cα RMSD of 0.40 Å (Fig. 2D, 2E; Extended Data Fig. 4C). At the same time, the Cap binding did not significantly affect the structure of Nsp10. We also compared the two structures of Nsp10/16/SAM determined at 297 K (this work PDB entry 7JIB) and 100 K (PDB entry 6W4H) (Cα RMSD: 0.35 Å) and observed significant differences: one loop from the Cap binding pocket is shifted approximately 1.4 Å in 297 K structures, which makes the Cap binding site more accessible (Extended Data Fig. 4D). Therefore, the 297 K structures of Nsp10/16 in a complex with SAM, Cap-0/SAM, and Cap-1/SAH, as determined by SSX, depict states important for molecular modeling studies and structure-based drug design.

To better understand Nsp16 activity from SARS-CoV-2, we compared it with NS5 2′-O MTase from Dengue virus (PDB entry 5DTO) (Fig. 3). We aligned the structure of Nsp16 with the N-terminal domain of NS5 that spans 298 residues from N-terminus. Both Nsp16 and NS5 2′-O MTases assume a Rossmann fold with a large β-sheet decorated with α-helices (Fig. 3B, D). MTases contain a central β-sheet that is composed of six parallel and one antiparallel strands. Despite low sequence identity (13.6%) the RMSD between the two MTases is reasonable (3.01 Å). Both enzymes possess a canonical 2′-O MTase catalytic tetrad Lys-Asp-Lys-Glu with the aspartic acid residue forming a hydrogen bond with a water molecule potentially relevant for catalysis (Fig. 3A, C). The SAM binding sites share high similarity but there are major differences in the mRNA Cap binding sites. The Mg^2+^ or Mn^2+^ ion is necessary for Nsp16 activity ^39^, however is not localized near the active site in the currently available structures of Nsp10/16 from SARS-CoV-1 and SARS-CoV-2. The Mg^2+^ ion in the N-terminal domain of NS5 2′-O MTase is coordinated by the phosphate oxygens of the Cap 5′ to 5′ triphosphate linker and three water molecules that bridge to three bases (A_1_-G2-U_3_) on the 5′-end.

**Figure 3.**
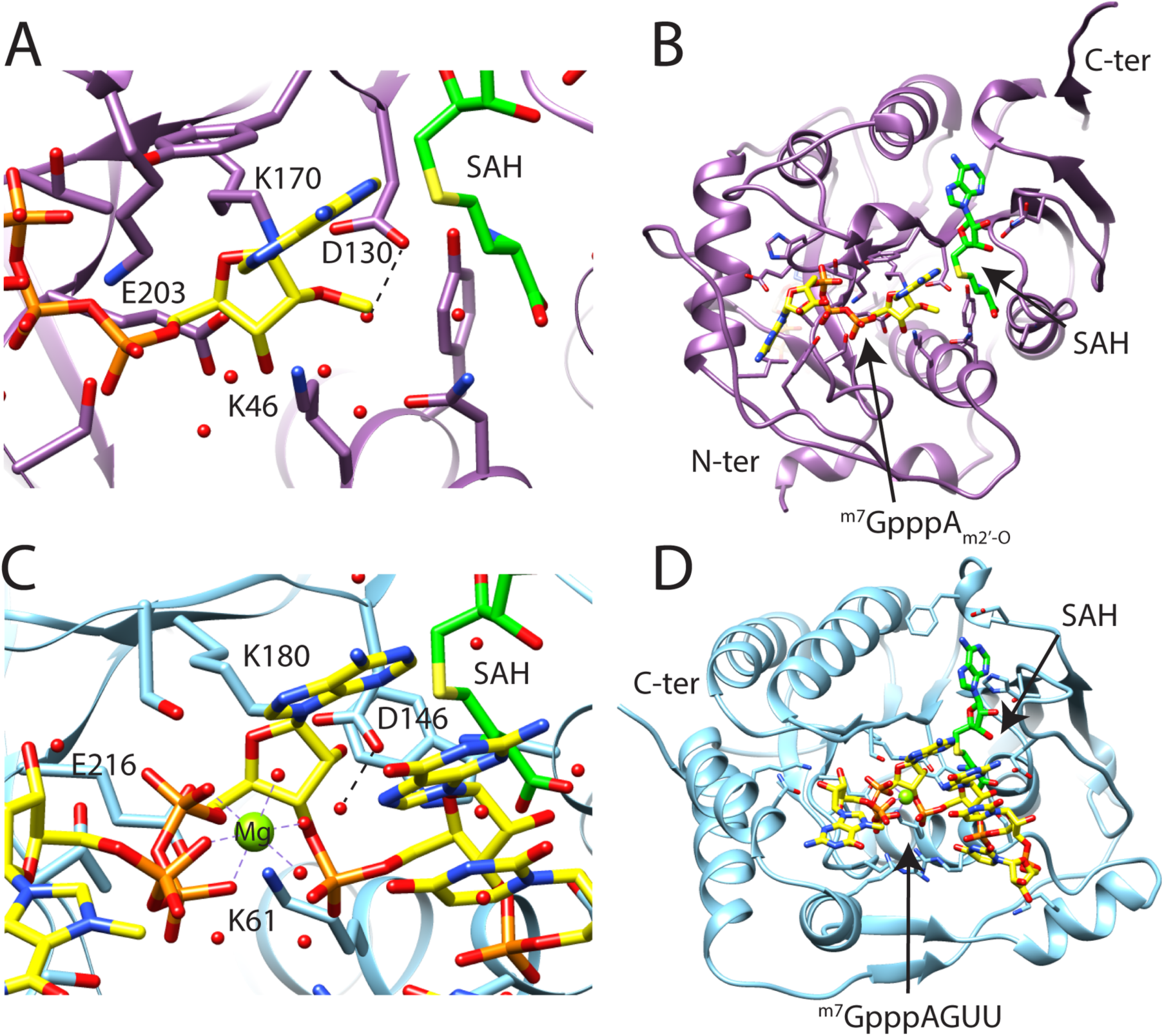
The Nsp16 from SARS-CoV-2 shares structural similarity with the N-terminal fragment of NS5 MTase from Dengue virus. Active site of Nsp16 from SARS-CoV-2 with catalytic tetrad Lys46-Asp130-Lys170-Glu203 (**A**). Structure of Nsp16 from SARS-CoV-2 in complex with ^m7^GpppA_m2′-O_ (Cap-1) and SAH (PDB entry 7JHE) (**B**). Active site of the N terminal fragment of NS5 from Dengue virus in a complex with ^m7^GpppAGUU and SAH shares conserved catalytic residues Lys61-Asp146-Lys180-Glu216 (**C**). The structure of the N terminal fragment of NS5 from Dengue virus (PDB entry 5DTO) (**D**).

### 2′-O methylation of the SARS-CoV-2 transcripts

The SARS-CoV-2 Nsp10/16 2′-O MTase complex provides a molecular arrangement for binding of the mRNA Cap-0 and subsequent methylation of the first transcribed nucleotide. The catalytic core is in the center of the Rossmann fold and it binds the SAM molecule in the deep, narrow grove that is buried inside the Nsp16 active site. We observed that Nsp16 had recruited SAM during expression in *E. coli* (PDB entry 6W61) ^20^. However, addition of 2 mM SAM to the crystallization solution improves crystallization efficiency, reduces crystallization time, and increases the number of crystals - three highly desirable traits for SSX batch crystallization. The SAM binding site is negatively charged and is formed by several Nsp16 residues: Asn43, Tyr47, Gly71, Ala72, Gly73, Ser 74, Gly81, Thr82, Asp99, Leu100, Asn101, Asp114, Cys115, Asp130, Met131, and Phe149 (Fig. 2, 4, Extended Data Fig. 5B). The SAM carboxylate moiety binds to the positively charged N-terminus of the third α-helix spanning Pro80-Trp88. This interaction provides additional electrostatic stabilization. Nsp16 directly interacts with Cap-1 through several residues that form a positively charged, elongated binding groove accommodating the mRNA Cap-0 (Fig. 2, 4, Extended Data 5A). The N7-metyl guanosine binding pocket is formed by Cys25, Asp26, Leu27, Tyr30, Thr172, Glu173, and Ser202. The 5′ to 5′ triphosphate bridge of the Cap is stabilized through interaction with Tyr30, Lys137, Thr172, His174, Ser201, Ser202, and Glu203. The first nucleotide of the mRNA Cap (adenosine in the presented structure) is bound through Lys46, Asp130, Tyr132, Pro134, Lys170, Asn198, and Glu203.

**Figure 4.**
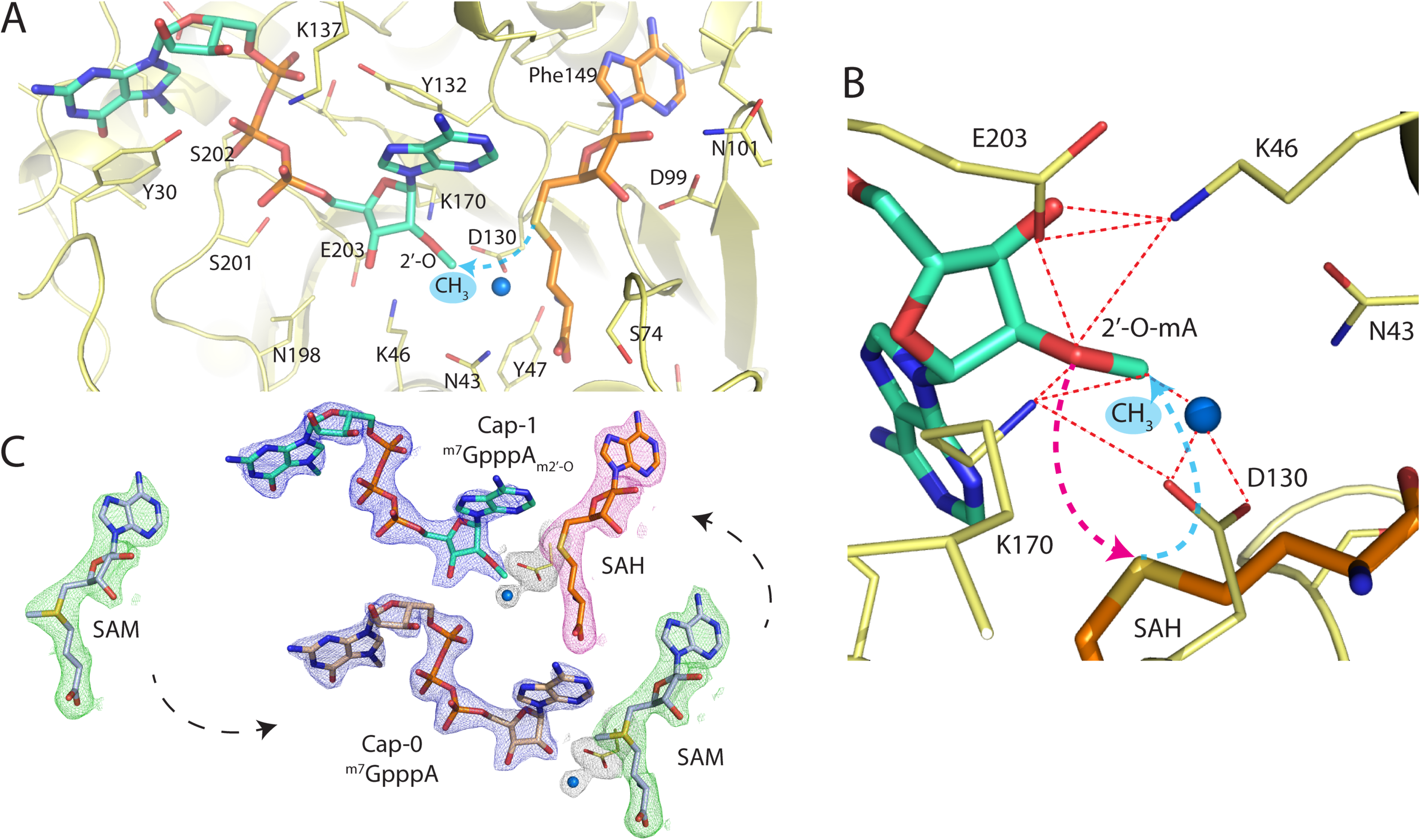
Methylation of the 2′-O-ribose of the ^m7^GpppA catalyzed by activity of the Nsp10/16 heterodimer from SARS-CoV-2. Active site of Nsp16 2′-O methyltransferase (yellow) in a complex with ^m7^GpppA_m2′-O_ (Cap-1) and SAH (**A**). Magnification of the active site depicts key residues Lys46-Asp130-Lys170-Glu203 essential for the 2′-O MTase activity (**B**). The 2Fo - Fc maps (σ = 1.2) contoured around ligands of the structures determined by SSX. The maps for ^m7^GpppA_m2′-O_ with SAH are colored on blue and red respectively. The map for the SAM structure is depicted as green (**C**).

SAM-dependent MTases share a conserved catalytic mechanism wherein the methyl group is transferred to the acceptor substrate via an S_N_2 reaction ^40^, which requires a linear alignment of the acceptor substrate (nucleophile), methyl group (electrophile), and the sulfur atom (leaving group) of the SAH product ^41^. It was proposed that Nsp16 2′-O MTases follow the same general mechanism ^42^. In these enzymes, the reaction is facilitated by the catalytic tetrad Lys-Asp-Lys-Glu, where Lys170 sandwiched between the two acidic residues (Asp130 and Glu203) serves as a proton abstractor (Fig. 4A, B). It was shown previously in biochemical studies that substitution to Ala of any residue of the 2′O-MTase catalytic tetrad results in an inactive enzyme ^10,43^. In SARS-CoV-2 Nsp16 the catalytic tetrad Lys46-Asp130-Lys170-Glu203, is superposing well with Lys61-D146-Lys180-Glu216 of the Dengue virus homolog (Fig. 3A, C). In the 3D context, Lys46 binds to Glu203 which then binds to Lys170 interacting with Asp130, which further links to the amino group of SAM. In this network, a proton can be transferred between multiple residues, occupying different sites potentially depending on the reaction state (Fig. 4). Lys170 is well-positioned to act as a general base deprotonating 2′-OH. Asp130 may serve multiple functions - as an acid deprotonating Lys170 (alternating with Glu203), and an anchoring point for the cofactor via interaction with its amino group, and as a stabilization for the sulfonium cation. However, the reaction does not occur in a presence of EDTA, as demonstrated by the ability to capture the Cap-0/SAM complex. It was reported previously using biochemical assays that the activity of Nsp10/16 2′-O MTase is magnesium dependent ^39^. Though, we did not observe any trace of metal near the active site in the 297 K structures reported in this work, and any other CoV-2 Nsp10/16 structures reported to date, despite the presence of magnesium in the crystallization buffer. We hypothesize that the magnesium ion (or other metal ion) can transiently bind to the active site possibly replacing one of the water molecules (for example water 38) and promoting formation of a reactive conformation by changing electrostatics and geometry of the catalytic residues to stimulate methylation reaction. Magnesium has a compact and tight coordination sphere with strict octahedral geometry and a typically short Mg—O distance of 2.08 Å ^44^. By coordinating 2′ oxygen and several water molecules magnesium could shortened the distance between 2′ oxygen and SAM methyl moiety thus promoting formation of transition state and methyl transfer. During the reaction a positively charged, sp2 planar transition state is formed and the methyl group inverts its stereochemistry.

After the methyl-transfer reaction is completed the product is released from the active site. This is consistent with the structure of Cap-1/SAH where several active site residues move, these include Tyr132, the entire α-helix spanning Pro134-Lys137, and Tyr30 on the opposite site (Fig. 4A, Extended Data Fig. 4C). All these residues are involved in interactions with Cap-0 and observed changes perhaps allow Cap-1 to leave. Interestingly, the SAM/SAH binding site remains virtually unchanged in all three structures (Extended Data Fig. 4), suggesting that the SAH exchange with SAM may require dissociation of Nsp10 that controls conformation of important loop Gly73-Gly77. Opening this loop may help SAH to leave and then a new SAM molecule can bind.

The biochemical assays show that optimal pH for MERS-CoV Nsp10/16 activity is approximately 8 - 8.5 ^39^. Additionally, the 2′-O MTase activity of Nsp10/16 from the MERS is significantly reduced at pH below 7. Assuming that Nsp10/16 2′-O MTase efficiency is pH dependent, we performed the 297 K SSX data collection using a buffer with pH 6.5 that could potentially slow down the reaction. Comparison of the 100 K and the 297 K structures revealed clear methylation only at 297 K, we hypothesize that Cap-1 could be sensitive to radiation damage and thus can be only observed in low x-ray dose experiments. Interestingly, the 297 K structure of Nsp10/16 obtained from a single crystal has a mixture of states with 30% of the Cap-1/SAH products and 70% of the Cap-0/SAM substrates. It is well known that radiation damage can impact redox systems. High x-ray dose could affect electron density around amino acid residues and lead to photoreduction of metalloproteins ^45,46,47^. These free radical reactions can occur in crystals under cryogenic conditions ^48^.

## CONCLUSIONS

Enzymes that catalyze transmethylation reactions using SAM as the methyl group donor have been described in many cellular processes involving nucleic acids, proteins, phospholipids, and small molecules ^49^. SARS-CoV-2 Nsp16 methyltransferase is one representative of this large family of proteins, which, together with Nsp10, participates in the post-transcriptional modification of the viral RNA. This multi-step process is a prerequisite to yield fully functional mRNA that (i) would not be recognized as foreign genetic element and, as such, be directed for degradation ^50^ and (ii) could undergo translation in the human host ^12,51^. 2′-O Methylation of the mRNA Cap facilitated by the Nsp10/16 is the last reaction in the mRNA assembly. Given the indispensable role of the RNA modification for the survival of the virus, the disturbance of this pathway, including the 2′-O methylation of Cap, is a field for development of inhibitors that affect SARS-CoV-2 replication. To that end, understating structure-function relationship of the enzymes catalyzing these process is critical.

Six months after the beginning of the COVID-19 outbreak we reported here the first structure for the SARS-CoV-2 Nsp10/16 complex revealing the post-reaction state with 2′-O methylated ribose of the Cap-1 bound to the protein. Despite several previous trials to visualize the product binding by traditional crystallography, no such state could be captured. Here we applied SSX as another attempt to fill the knowledge gaps in the 2′-O methylation process. The SSX experiments on the Nsp10/16 crystals revealed three states: with SAM bound, with SAM and Cap-0 prior reaction occurring, and, finally with Cap-1 and SAH after methyl transfer. These experiments were done at 297 K under low x-ray radiation dose to the crystal. In comparison, the 297 K single crystal structure shows methylation of the 2′-O-ribose of the Cap-0, but with partial occupancy. Interestingly, the methylation was not observed previously in any of the Nsp10/16 structures determined at 100 K ^20,24,42,52,53^. Is this reaction sensitive to radiation damage or free radical effects? It seems that the most elusive may be the formation of the metal-dependent state resulting in establishing the electrostatic/geometric conditions prior activation or destabilization of transition state as there is no depletion of the methyl group on SAM in any reported structures. This clearly needs further investigation because SSX low radiation dose may be advantageous in helping reveal subtle and diverse chemical transformations in enzymes that otherwise may be degraded during x-ray or electron diffraction experiments. Therefore, the structure of Nsp10/16 with Cap-1 presented here provides unique information for understanding the mechanism that allows SARS-CoV-2 to mimic mature eukaryotic mRNA and escape recognition by human innate immune response.

The recent developments in micro focusing x-ray beams at synchrotron light sources and improvement in sample delivery technology, data collection, detectors, and computing will allow rapid determination of new structures using SSX and revealing significant biological information. The further development of SSX and implementation of time-resolved SSX crystallography is an approach that could visualize chemical processes and protein molecular dynamics - such as of the transfer of the methyl group catalyzed by Nsp10/16 2′O-MTase from SARS-CoV-2. Thus far the Cap-1 was only observed in the structure obtained using SSX. For diffraction experiments conducted at 297 K, SSX provides significant advantages over data collected using a single crystal as it considerably increases resolution, use less sample in comparison with a liquid jet delivery system and reduces levels of x-ray dose. Studies of other enzymes can also significantly benefit from using this approach.

## METHODS

### Crystallization of Nsp10/16 complex

The recombinant proteins Nsp10 and Nsp16 from SARS-CoV-2 were expressed in *E. coli* and purified as described by Rosas-Lemus ^20^. The freshly purified Nsp10 and Nsp16 were mixed in a 1:1 molar ratio to the final concentration of the complex as 2 mg/ml. Subsequently, the Nsp10/16 sample was dialyzed overnight at 297 K in a buffer containing 10 mM Tris-HCl, 150 mM NaCl, 2 mM MgCl_2_, 1 mM TCEP, 5% w/v glycerol. The Nsp10/16 heterodimer was supplemented with 2 mM SAM (Sigma Aldrich) and concentrated at 21°C to 4.0 mg/ml using a 30 kDa cut-off centrifugal concentrator Amicon Ultra-15 (Millipore). Two criteria for preliminary selection for batch crystallization were getting crystals with the highest possible symmetry as well as having one Nsp10/16 heterodimer in the asymmetric unit. Such conditions were met by crystals with hexagonal space group P3_1_21. The complex was initially crystallized in a buffer 0.1 M MES pH 6.5 and 0.6 M sodium fluoride from Anions Suite crystallization screen (Qiagen). After optimization of crystallization conditions, we used 0.1 M MES pH 6.5, 0.9 M NaF for batch crystallizations. For each batch of crystals 100 µl of the Nsp10/16 complex was mixed with 100 µl of the precipitant buffer and crystallization was performed in 500 µl Eppendorf polypropylene tube. These volumes allowed us to get the number of crystals suitable for data collection from one chip. We have prepared several batches using same conditions to get reproducible data collection and have the ability to merge data from multiple chips. The crystals grew to optimum sizes in 10 days at 21°C before the SSX data collection. To obtain the structure of Nsp10/16/Cap-0/SAM, the crystal batch was supplemented with 1 mM EDTA two days before data collection.

### Preparation of the sample for data collection

The crystals from batch crystallization were centrifuged at 100 RCF for 2 minutes at 297 K. The excess solution was removed with a pipette, until 15 µl was left. Crystals sedimented on the walls of Eppendorf polypropylene tube were gently resuspended using 20 µl pipette tips with a cut end to increase the diameter of the tip to minimize mechanical damage to crystals. Then 1.5 µl of 10 mM stock of the ^m7^GpppA (S1405L, New England Biolabs) was added to crystal slurry and the mix was loaded on a 60 µm grid made from nylon (NY6004700, Millipore) which was placed on a 6 µm layer of mylar polyester film. The nylon mesh was covered on the top with a second layer of 6 µm mylar and sealed with the *ALEX* magnetic holder, such that sample is hermetically sealed from the outside environment. Data collection was started approximately 20 minutes after assembly of the chip.

### Serial crystallography data collection

We collected SSX data for the Nsp10/16 crystals at the 19-ID beamline at the Advanced Photon Source using the fixed-target SSX (*ALEX)* mesh-holder developed at the Structural Biology Center (SBC) as depicted in Figure 1A and Extended Data Fig. 1. A crystal slurry is deposited on a nylon mesh, which immobilizes the crystals; they are then encapsulated between two polyester films ^36^. The rod-shape crystals grew without seeding and the average crystal size was 120×25×25 µm (Extended Data Fig. 2A). Serial data collection was collected using three SmarAct SLC-17 stages configured in an XYZ geometry, with each having sufficient movement range to cover the sample area of the specially designed *ALEX* holder (patent application serial #16/903,601). The beamline was configured at an energy of 12,662 eV, with collimator slit sizes set to 75 x 75 µm, and step size (distance between exposures) of 50 µm: overlapping exposed area to maximize crystal hits. The five mesh-covered samples used a grid of 170 steps in the x-direction (columns) by 210 steps in the y-direction (rows), covering a total area of approximately 8.5 x 10.5 mm. The number of steps and the resulting area varied slightly per sample depending on the chip mount position or possible false-starts. Extended Data Table 1 contains details of number of chips and detector distances used for data collection for Nsp10/16 SSX structures.

*Crispy*, the data acquisition GUI for serial collection at sector 19, allows for quick alignment and acts as a source of information for downstream processing. Metadata in the form of JSON files and beamline/collection strategy parameters are input and passed into the system before collection. These parameters include grid dimensions, detector distance/resolution, unit cell dimension, protein PDB coordinates, and a handful of others.

### Serial crystallography data processing

The *Kanzus* pipeline orchestrates SSX data acquisition, analysis, cataloging, and publishes processing metrics. *Kanzus* uses a cloud-hosted research automation system called Globus Automate to manage these multi-step data “flows” ^54^. The first phase of the pipeline is integrated with the APS Data Management System at the beamline, which deposits each newly acquired image into an Globus-accessible storage system at the APS. As new images are acquired, Globus Automate “flows” are launched to process them as follow: 1) moves new files from APS to Theta by using the Globus Transfer service ^55^; 2) performs DIALS *stills_process* ^56^ on batches of 256 images by using funcX ^57^, a function-as-a-service computation system (funcX uses Parsl ^58^ to abstract and acquire nodes on Theta as needed, and dispatches tasks to available nodes); 3) extracts metadata from files regarding identified diffractions, and generates visualizations (funcX) showing the locations of positive hits on the mesh; and 4) publishes raw data, metadata, and visualizations to a portal on the ALCF Petrel data system ^59^. The result of this automated process is an indexed, searchable data collection that provides full traceability from data acquisition to processed data, and that can be used to inspect and update the running experiment.

Images were collected at 7 Hz, meaning that a 256-image batch totaling 1.56 GB, was generated every 35 seconds. The data transfers to ALCF ran at up to 700 MB/s via Globus, and 30 Theta nodes processed images by using DIALS *stills_process* at 22 images per second in steady-state. As experimental configuration values were refined, reprocessing tasks were submitted as required. FuncX managed these tasks by expanding the number of Theta nodes being used to a maximum of 250, which enabled a processing rate of greater than 200 images a second. Successfully processed images with diffraction-produced integration files were returned to the beamline computers and later refined and merged using PRIME ^60^.

### Structure solution and refinement

The crystal structures of the Nsp10/16 complex were solved by molecular replacement using MolRep ^61^ from the CCP4 package. The structures were refined by multiple cycles in REFMAC v. 5.8.0258 ^62^ followed by manual edition of the model using Coot ^63^. Ligands: Cap-1, Cap-0, ^m7^Gppp, ^m7^Gpp, SAM, SAH, and Zn^2+^ were manually placed in to electron density at the beginning of the refinements. The stereochemistry of structures were analyzed using MolProbity and the Ramachandran plot. The atomic coordinates and structure factors for Nsp10/16 structures determined with SSX were deposited to PDB with assigned accession codes: 6XKM – SAM, 7JPE – Cap-0/SAM, 7JHE – Cap-1/SAH. Nsp10/16 complex with mixture of Cap-0/Cap-1 and SAH/SAM determined from single crystal at 297 K has PDB entry 7JIB. Figures were prepared with PyMol and UCSF Chimera ^64^.

## Author contributions

Wilamowski M., Sherrell D.A., Minasov G., Shuvalova L., Foster I.T, Satchell K.J.F. and Joachimiak A. designed the research. Wilamowski M., Shuvalova L., Rosas-Lemus M., Maltseva N. and Jedrzejczak R. purified Nsp10/16 complex and crystallized protein. Sherrell D.A., Lavens A., and Chard R. performed serial crystallography data collection. Foster I.T. contributed to the distributed processing pipeline. Wilamowski M., Sherrell D.A., Minasov G., Kim Y., Saint N., and Chard R analyzed serial crystallography data. Wilamowski M., Sherrell D.A., Minasov G., Lavens A., Michalska, K., and Joachimiak A. wrote the manuscript.

## EXTENDED DATA

**Extended Data Table 1.** Data collection and processing statistics for Nsp10/16 297 K structures. Comparison of data collected using fixed-target serial crystallography and using single crystal mounted in capillary.

|  | **Nsp10/16**<br>**SAM** | **Nsp10/16**<br>**Cap-1 + SAH** | **Nsp10/16**<br>**Cap-0 + SAM** | **Nsp10/16**<br>**Cap-0/Cap-1 +**<br>**SAM/SAH** |
| --- | --- | --- | --- | --- |
| **Data Collection** | fixed-target SSX | fixed-target SSX | fixed-target SSX | single crystal |
| Crystal holder | 60 μm nylon mesh<br>between 2 layers of<br>mylar | 60 μm nylon mesh<br>between 2 layers of<br>mylar | 60 μm nylon mesh<br>between 2 layers of<br>mylar | MiTeGen microRT<br>capillary |
| Crystal size (μm) | 120x25x25 | 120x25x25 | 120x25x25 | 250x50x50 |
| Diffraction source | APS beamline 19-ID | APS beamline 19-ID | APS beamline 19-ID | APS beamline 19-ID |
| Wavelength ($\text{\AA}$) | 0.9792 | 0.9792 | 0.9792 | 0.9792 |
| Temperature (K) | 297 | 297 | 297 | 297 |
| Detector | PILATUS3 X 6M | PILATUS3 X 6M | PILATUS3 X 6M | PILATUS3 X 6M |
| Beam size (μm) | 75 x 75 | 75 x 75 | 75 x 75 | 50 x 50 |
| Exposure time per image (s) | 0.05 | 0.05 | 0.05 | 0.2 |
| Beam attenuation | 1 | 1 | 1 | 2.2 |
| Crystal – detector distance (mm) | 540 (1<sup>st</sup> chip)<br>319 (2<sup>nd</sup> chip) | 319 (1<sup>st</sup> and 2<sup>nd</sup> chip)<br>446 (3<sup>rd</sup> chip) | 422 (1<sup>st</sup> and 2<sup>nd</sup>) | 470 |
| Number of chips | 2 | 3 | 2 | - |
| Number of collected images | 73100 | 105740 | 74120 | 180 (90°) |
| Number of indexed images | 10990 | 7725 | 5734 | 180 |
| Indexing hit rate (%) | 15.0 | 7.3 | 7.8 | - |
| Diffraction images resolution lower than 3.5$\text{\AA}$ | 1622 | 1654 | 976 | - |
| Diffraction images < 90 reflections | 635 | 428 | 187 | - |
| Diffraction images with deviation from the<br>crystal unit cell volume > 4.5% | 2009 | 1831 | 1362 | - |
| Number of images after rejection of outliers | 7951 | 5038 | 4704 | - |
| Images with indexed single lattice | 6985 | 4522 | 4098 | 180 |
| Images with indexed two lattices | 900 | 488 | 555 | - |
| Images with indexed three lattices | 66 | 28 | 51 | - |
| **Lattices merged for structure determination** | 7267 | 4548 | 4274 | 180 |
| **Total time of data collection (min)** | 180 | 270 | 180 | < 1 |
| **Dose of X-ray per crystal (MGy)** | 0.12 | 0.12 | 0.12 | 4.97 |

**Extended Data Table 2.** Crystallographic data and refinement statistics.

| **Data Collection** | **Nsp10/16**<br>**SAM**<br>fixed-target SSX | **Nsp10/16**<br>**Cap-1 + SAH**<br>fixed-target SSX | **Nsp10/16**<br>**Cap-0 + SAM**<br>fixed-target SSX | **Nsp10/16**<br>**Cap-0/Cap-1 +**<br>**SAM/SAH**<br>single crystal in capillary |
| --- | --- | --- | --- | --- |
| Space group | *P3<sub>1</sub>21* | *P3<sub>1</sub>21* | *P3<sub>1</sub>21* | *P3<sub>1</sub>21* |
| a, b, c (Å) | 170.41, 170.41, 52.63 | 170.86, 170.86, 52.80 | 170.93, 170.93, 52.79 | 170.83, 170.83, 52.74 |
| $\alpha, \beta, \gamma$ (°) | 90.0, 90.0, 120.0 | 90.0, 90.0, 120.0 | 90.0, 90.0, 120.0 | 90.0, 90.0, 120.0 |
| Resolution range (Å)<sup>a</sup> | 49.57 – 2.25 (2.29 – 2.25) | 49.73 – 2.25 (2.29 – 2.25) | 49.72 – 2.18 (2.22 – 2.18) | 50.00 – 2.65 (2.70 – 2.65) |
| No. of unique reflections | 41,770 (2,090)<sup>a</sup> | 42,145 (2,126)<sup>a</sup> | 46,273 (2,290)<sup>a</sup> | 25,799 (1,263)<sup>a</sup> |
| Completeness (%) | 100.0 (99.7) | 100.0 (99.8) | 100.0 (99.7) | 99.6 (99.8) |
| Data redundancy | 82.41 (10.24) | 42.94 (9.28) | 50.51 (11.42) | 5.2 (4.9) |
| R<sub>split</sub> (%)<sup>b</sup> | 18.75 (96.45) | 26.96 (87.30) | 23.44 (98.03) | - |
| R<sub>p.i.m.</sub> (%)<sup>c</sup> | - | - | - | 0.071 (0.507) |
| CC/1/2<sup>d</sup> | 0.968 (0.449) | 0.930 (0.473) | 0.950 (0.398) | 0.981 (0.537) |
| $\langle I/\sigma(I) \rangle$ | 2.68 (0.58) | 2.45 (0.66) | 2.55 (0.53) | 15.0 (1.87) |
| Wilson B factor | 23.2 | 21.4 | 22.8 | 49.9 |
| **Structure Determination** |  |  |  |  |
| MR initial model (PDB ID) | 6W4H | 6WQ3 | 6WQ3 | 6WQ3 |
| **Refinement** |  |  |  |  |
| Resolution range (Å) | 49.57 – 2.25 (2.31 – 2.25) | 49.73 – 2.25 (2.31 – 2.25) | 49.72 – 2.18 (2.22 – 2.18) | 29.59 – 2.65 (2.72 – 2.65) |
| Completeness (%) | 99.9 (98.8) | 99.9 (99.5) | 99.97 (99.74) | 99.0 (98.9) |
| No. of unique reflections | 39,686 (2,893) | 40,062 (2,962) | 46273 (2290) | 24,366 (1,751) |
| R<sub>work</sub>/R<sub>free</sub><sup>e</sup> (%) | 20.65/21.26 (36.3/33.4) | 22.31/24.85 (37.4/36.0) | 21.70/23.70 (37.9/34.1) | 15.49/18.09 (31.2/30.2) |
| Protein chains/atoms | 2/3,204 | 2/3,210 | 2/3,210 | 2/3,244 |
| Ligand/Solvent atoms | 30/129 | 114/160 | 109/131 | 189/141 |
| Mean temperature factor (Å<sup>2</sup>) | 37.6 | 37.1 | 40.0 | 55.9 |
| **Coordinate Deviations** |  |  |  |  |
| R.m.s.d. bonds (Å) | 0.004 | 0.005 | 0.005 | 0.004 |
| R.m.s.d. angles (°) | 1.213 | 1.363 | 1.351 | 1.291 |
| **Ramachandran plot<sup>f</sup>** |  |  |  |  |
| Favored (%) | 96.0 | 95.0 | 97.0 | 95.0 |
| Allowed (%) | 4.0 | 5.0 | 3.0 | 5.0 |
| Outside allowed (%) | 0.0 | 0.0 | 0.0 | 0.0 |
| **PDB Accession Code** | **6XKM** | **7JHE** | **7JPE** | **7JIB** |

<sup>a</sup>Values in parentheses correspond to the highest resolution shell.
<sup>b</sup>R<sub>split</sub> as defined by White<sup>65</sup>.
<sup>c</sup>R<sub>p.i.m.</sub> as defined by Weiss<sup>66</sup>.
<sup>d</sup>CC/1/2 as defined by Karplus and Diederichs<sup>67</sup>.
<sup>e</sup>R = $\Sigma |F_o| - |F_c| / \Sigma |F_o|$ for all reflections, where F<sub>o</sub> and F<sub>c</sub> are observed and calculated structure factors, respectively. R<sub>free</sub> is calculated analogously for the test reflections, randomly selected and excluded from the refinement.
<sup>f</sup>As defined by Molprobity<sup>68</sup>.

**Extended Data Figure 1.**
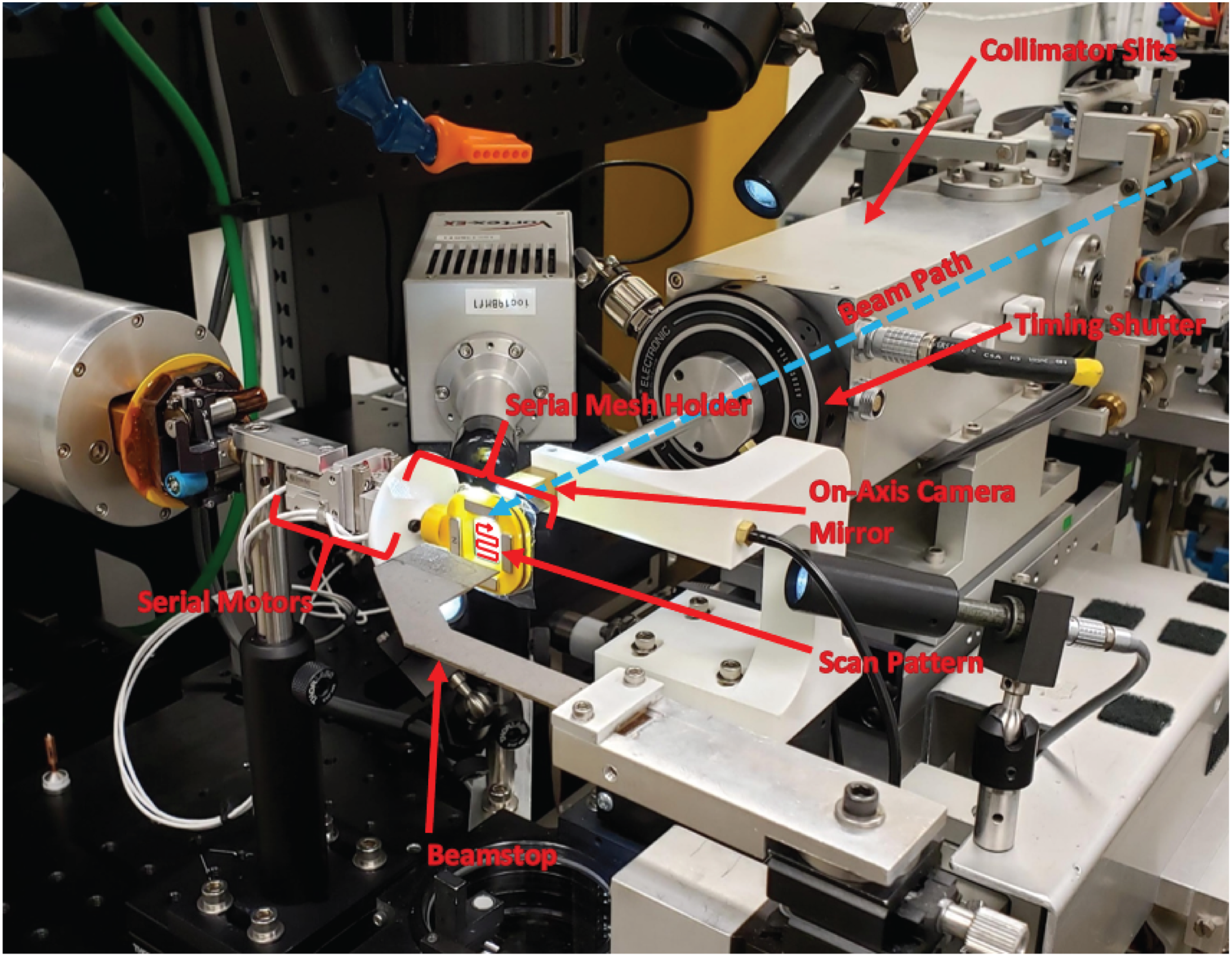
Fixed-target SSX system at Structural Biology Center beamline 19-ID at APS. The serial crystallography chip holds nylon mesh between the two layers of mylar.

**Extended Data Figure 2.**
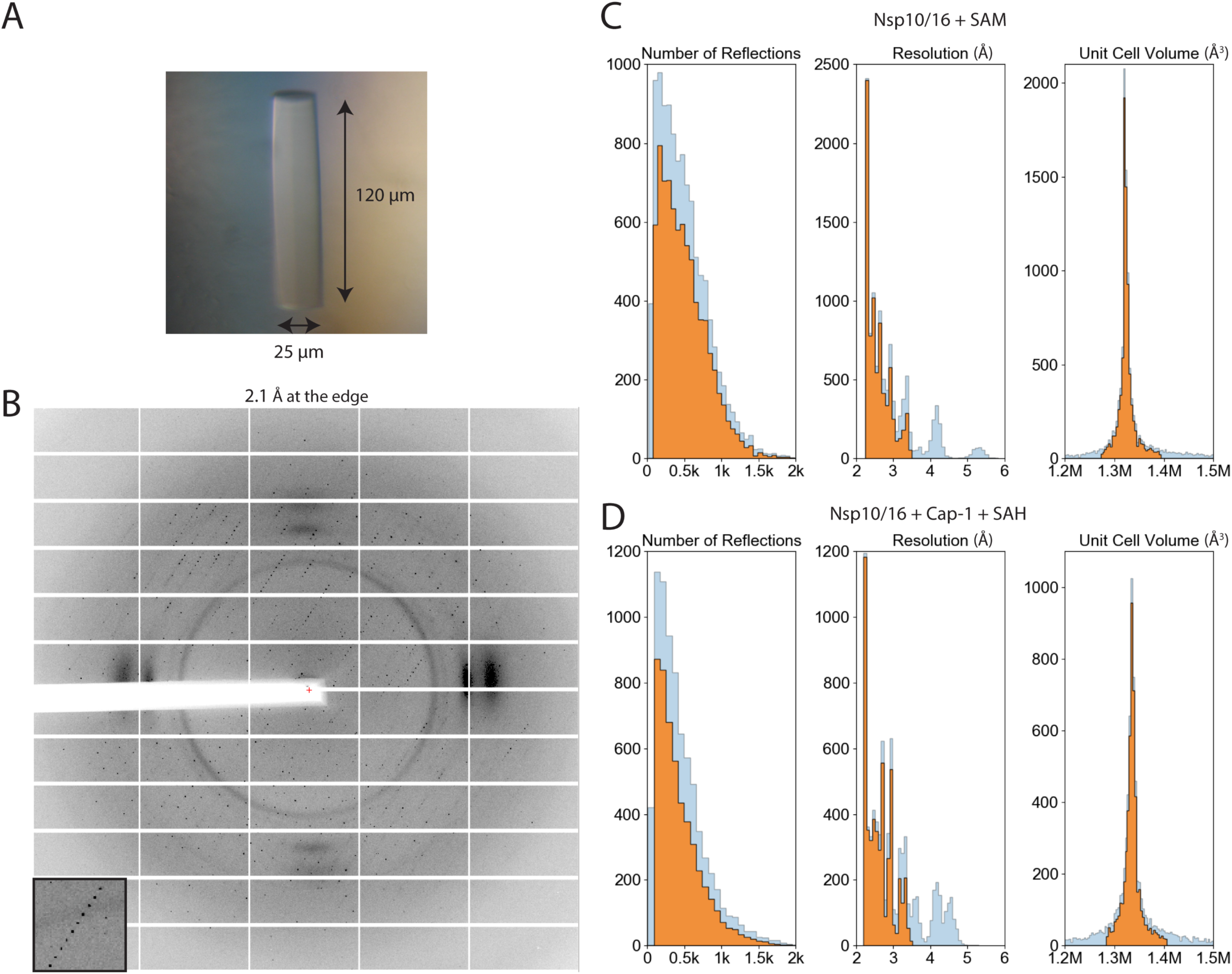
Processing of SSX data for Nsp10/16. Representative crystal of Nsp10/16 used for 297 K SSX data collection (**A**). Diffraction image of one of the Nsp10/16 crystals obtained during SSX data collection, detector edge at 2.1 Å. The fiber diffraction pattern visible at ~ 4 Å originates from the nylon mesh scattering. (**B**). Processing of SSX data with DIALS, histograms illustrates number of reflections, resolution and unit cell volume obtained for integrated images. Depicted integration statistics for Nsp10/16/SAM (**C**) and Nsp10/16/Cap-1/SAH (**D**). The blue bars show data before rejections of outliers (unit cell deviation > 4.5%, resolution < 3.5 Å, number of reflections < 90), orange bars depict data used for scaling and refinement of the Nsp10/16 structures.

**Extended Data Figure 3.**
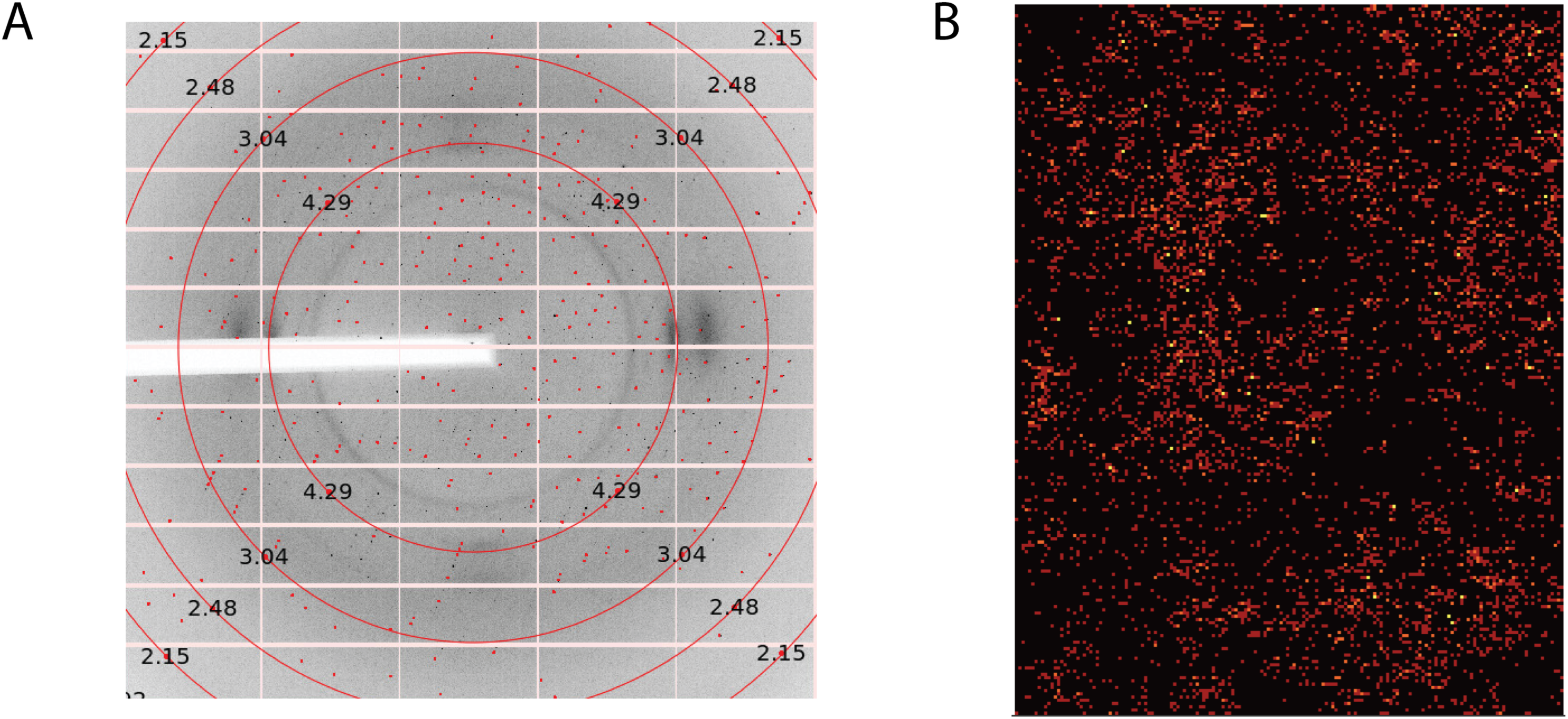
Fixed-target serial crystallography data collection for Nsp10/16 crystals. Diffraction image of Nsp10/16 crystals obtained from fixed-target SSX, image processed with DIALS viewer, red dots represents indexed crystal reflections (**A**). Hit map illustrates images integrated using DIALS, red dots – single lattice, orange dots – two lattices, yellow dots – three lattices determined on integrated crystal diffraction image (**B**).

**Extended Data Figure 4.**
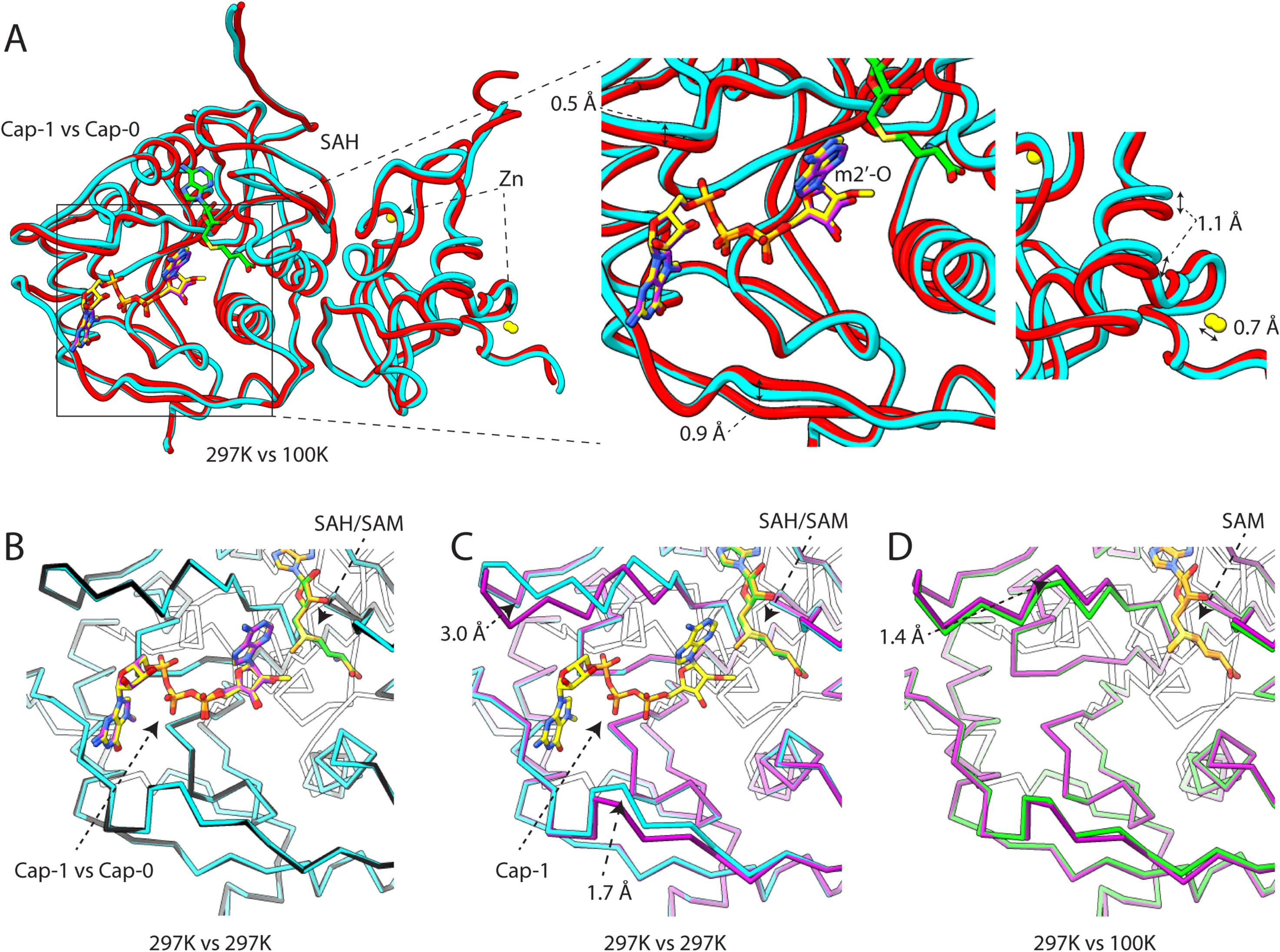
Comparison of 297 K crystal structures of the Nsp10/16 with structures determined at 100 K. Crystal structure of the Nsp10/16 complex with Cap-1 and SAH (blue – 7JHE) determined using SSX superimposed with Nsp10/16 complex with Cap-0 and SAH (red - 6WQ3) determined at 100 K (**A**). Comparison of 297 K crystal structures of Nsp10/16/Cap-1/SAH (blue – 7JHE) superimposed to structure of Nsp10/16/Cap-0/SAM (black – 7JPE) (**B**). Comparison of 297 K crystal structures of the Nsp10/16/Cap-1/SAH (blue – 7JHE) superimposed to structure of Nsp10/16/SAM (pink – 6XKM) (**C**). Comparison of the Nsp10/16 crystal structures with SAM determined respectively using SSX (pink – 6XKM) and standard data collection at 100 K (white - 6W4H) (**D**).

**Extended Data Figure 5.**
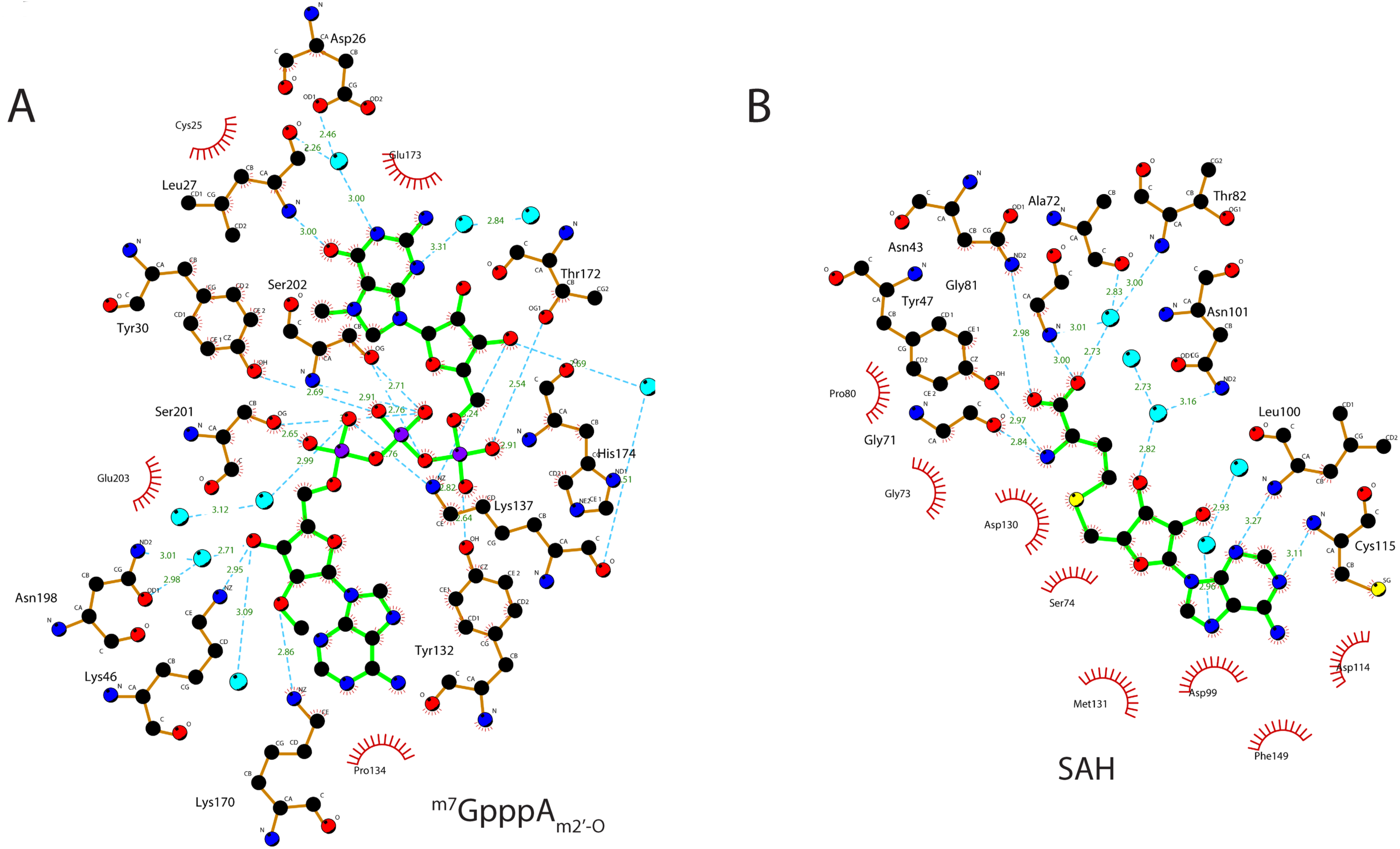
Hydrogen bonds and hydrophobic interactions that stabilize binding of ^m7^GpppA_m2′-O_ and SAH by Nsp10/16 2′-O MTase from SARS-CoV-2. Schematic diagrams of Nsp10/16 interactions with ligand made using LIGPLOT v.4.5.3 software ^69^.

## REFERENCES

1. Stadler, K. et al. SARS — beginning to understand a new virus. Nat. Rev. Microbiol. 1, 209–218 (2003).

2. Huang, C. et al. Clinical features of patients infected with 2019 novel coronavirus in Wuhan, China. Lancet 395, 497–506 (2020).

3. Chan-Yeung, M. & Xu, R. H. SARS: Epidemiology. Respirology vol. 8 (2003).

4. WHO | Middle East respiratory syndrome coronavirus (MERS-CoV). WHO (2020).

5. COVID-19 Map - Johns Hopkins Coronavirus Resource Center. https://coronavirus.jhu.edu/map.html.

6. Wu, A. et al. Genome Composition and Divergence of the Novel Coronavirus (2019-nCoV) Originating in China. Cell Host Microbe 27, 325–328 (2020).

7. Nagarajan, V. K., Jones, C. I., Newbury, S. F. & Green, P. J. XRN 5’→3’ exoribonucleases: Structure, mechanisms and functions. Biochimica et Biophysica Acta - Gene Regulatory Mechanisms vol. 1829 590–603 (2013).

8. Ramanathan, A., Robb, G. B. & Chan, S. H. mRNA capping: Biological functions and applications. Nucleic Acids Research (2016) doi:10.1093/nar/gkw551.

9. Ferron, F., Decroly, E., Selisko, B. & Canard, B. The viral RNA capping machinery as a target for antiviral drugs. Antiviral Research vol. 96 21–31 (2012).

10. Bouvet, M. et al. In Vitro Reconstitution of SARS-Coronavirus mRNA Cap Methylation. PLoS Pathog. 6, e1000863 (2010).

11. Werner, M. et al. 2′-O-ribose methylation of cap2 in human: Function and evolution in a horizontally mobile family. Nucleic Acids Res. (2011) doi:10.1093/nar/gkr038.

12. Muthukrishnan, S., Moss, B., Cooper, J. A. & Maxwell, E. S. Influence of 5’-terminal cap structure on the initiation of translation of vaccinia virus mRNA. J. Biol. Chem. (1978).

13. Hyde, J. L. & Diamond, M. S. Innate immune restriction and antagonism of viral RNA lacking 2’-O methylation. Virology vols 479-480 66–74 (2015).

14. Züst, R. et al. Ribose 2’-O-methylation provides a molecular signature for the distinction of self and non-self mRNA dependent on the RNA sensor Mda5. Nat. Immunol. 12, 137–143 (2011).

15. Menachery, V. D., Debbink, K. & Baric, R. S. Coronavirus non-structural protein 16: Evasion, attenuation, and possible treatments. Virus Research vol. 194 191–199 (2014).

16. Schuberth-Wagner, C. et al. A Conserved Histidine in the RNA Sensor RIG-I Controls Immune Tolerance to N1-2’O-Methylated Self RNA Immunity 43, 41–51 (2015).

17. Daffis, S. et al. 2′-O methylation of the viral mRNA cap evades host restriction by IFIT family members. Nature 468, 452–456 (2010).

18. Byszewska, M., Smietański, M., Purta, E. & Bujnicki, J. M. RNA methyltransferases involved in 5′ cap biosynthesis. RNA Biology vol. 11 1597–1607 (2014).

19. Decroly, E. et al. Coronavirus Nonstructural Protein 16 Is a Cap-0 Binding Enzyme Possessing (Nucleoside-2′O)-Methyltransferase Activity. J. Virol. 82, 8071–8084 (2008).

20. Rosas-Lemus, M. et al. The crystal structure of nsp10-nsp16 heterodimer from SARS-CoV-2 in complex with S-adenosylmethionine. bioRxiv Prepr. Serv. Biol. 2020.04.17.047498 (2020) doi:10.1101/2020.04.17.047498.

21. Bouvet, M. et al. Coronavirus Nsp10, a critical co-factor for activation of multiple replicative enzymes. J. Biol. Chem. 289, 25783–25796 (2014).

22. Bujnicki, J. M. & Rychlewski, L. In silico identification, structure prediction and phylogenetic analysis of the 2-O-ribose (cap 1) methyltransferase domain in the large structural protein of ssRNA negative-strand viruses. Protein Engineering vol. 15 http://bioinfo.pl/meta/ (2002).

23. Feder, M., Pas, J., Wyrwicz, L. S. & Bujnicki, J. M. Molecular phylogenetics of the RrmJ/fibrillarin superfamily of ribose 2′-O-methyltransferases. Gene 302, 129–138 (2003).

24. Chen, Y. et al. Biochemical and structural insights into the mechanisms of sars coronavirus RNA ribose 2′-O-methylation by nsp16/nsp10 protein complex. PLoS Pathog. 7, (2011).

25. Menachery, V. D. et al. Attenuation and Restoration of Severe Acute Respiratory Syndrome Coronavirus Mutant Lacking 2’-O-Methyltransferase Activity. J. Virol. 88, 4251–4264 (2014).

26. Barends, T. R. M. et al. Direct observation of ultrafast collective motions in CO myoglobin upon ligand dissociation. Science (80-.). 350, 445–450 (2015).

27. Olmos, J. L. et al. Enzyme intermediates captured ‘on the fly’ by mix-and-inject serial crystallography. BMC Biol. 16, (2018).

28. Weinert, T. et al. Proton uptake mechanism in bacteriorhodopsin captured by serial synchrotron crystallography. Science (80-.). 364, 61–65 (2019).

29. Chapman, H. N. et al. Femtosecond X-ray protein nanocrystallography. Nature 470, 73–78 (2011).

30. Boutet, S. et al. High-Resolution Protein Structure Determination by Serial Femtosecond Crystallography. Science (80-.). 337, 362–364 (2012).

31. Weinert, T. et al. Serial millisecond crystallography for routine room-temperature structure determination at synchrotrons. Nat. Commun. 8, (2017).

32. Weierstall, U. Liquid sample delivery techniques for serial femtosecond crystallography. Philosophical Transactions of the Royal Society B: Biological Sciences vol. 369 (2014).

33. Martin-Garcia, J. M. et al. High-viscosity injector-based pink-beam serial crystallography of microcrystals at a synchrotron radiation source. IUCrJ 6, 412–425 (2019).

34. Sherrell, D. A. et al. A modular and compact portable mini-endstation for high-precision, high-speed fixed target serial crystallography at FEL and synchrotron sources. J. Synchrotron Radiat. 22, 1372–1378 (2015).

35. Meents, A. et al. Pink-beam serial crystallography. Nat. Commun. 8, (2017).

36. Lee, D. et al. Nylon mesh-based sample holder for fixed-target serial femtosecond crystallography. Sci. Rep. 9, 1–9 (2019).

37. Zeldin, O. B., Gerstel, M. & Garman, E. F. RADDOSE-3D: Time- and space-resolved modelling of dose in macromolecular crystallography. J. Appl. Crystallogr. 46, 1225–1230 (2013).

38. Wang, Y. et al. Coronavirus nsp10/nsp16 Methyltransferase Can Be Targeted by nsp10-Derived Peptide In Vitro and In Vivo To Reduce Replication and Pathogenesis. J. Virol. 89, 8416–8427 (2015).

39. Aouadi, W. et al. Binding of the Methyl Donor S-Adenosyl-l-Methionine to Middle East Respiratory Syndrome Coronavirus 2′-O-Methyltransferase nsp16 Promotes Recruitment of the Allosteric Activator nsp10. J. Virol. 91, (2017).

40. Komoto, J. et al. Crystal structure of Guanidinoacetate methyltransferase from rat liver: A model structure of protein arginine methyltransferase. J. Mol. Biol. 320, 223–235 (2002).

41. Zhang, J., Kulik, H. J., Martinez, T. J. & Klinman, J. P. Mediation of donor-acceptor distance in an enzymatic methyl transfer reaction. Proc. Natl. Acad. Sci. U. S. A. 112, 7954–7959 (2015).

42. Decroly, E. et al. Crystal structure and functional analysis of the SARS-coronavirus RNA cap 2′-o-methyltransferase nsp10/nsp16 complex. PLoS Pathog. 7, (2011).

43. Martin, B. et al. The methyltransferase domain of the Sudan ebolavirus L protein specifically targets internal adenosines of RNA substrates, in addition to the cap structure. Nucleic Acids Res. (2018) doi:10.1093/nar/gky637.

44. Zheng, H. et al. CheckMyMetal: A macromolecular metal-binding validation tool. Acta Crystallogr. Sect. D Struct. Biol. 73, 223–233 (2017).

45. Ebrahim, A. et al. Dose-resolved serial synchrotron and XFEL structures of radiation-sensitive metalloproteins. IUCrJ 6, 543–551 (2019).

46. Fukuda, Y. et al. Redox-coupled proton transfer mechanism in nitrite reductase revealed by femtosecond crystallography. Proc. Natl. Acad. Sci. U. S. A. 113, 2928–2933 (2016).

47. Schlichting, I. et al. The catalytic pathway of cytochrome P450cam at atomic resolution. Science (80-.). 287, 1615–1622 (2000).

48. Petrova, T. et al. X-Ray-Radiation-Induced Cooperative Atomic Movements in Protein. J. Mol. Biol. 387, 1092–1105 (2009).

49. Struck, A. W., Thompson, M. L., Wong, L. S. & Micklefield, J. S-Adenosyl-Methionine-Dependent Methyltransferases: Highly Versatile Enzymes in Biocatalysis, Biosynthesis and Other Biotechnological Applications. ChemBioChem vol. 13 2642–2655 (2012).

50. Devarkar, S. C. et al. Structural basis for m7G recognition and 2′-O-methyl discrimination in capped RNAs by the innate immune receptor RIG-I. Proc. Natl. Acad. Sci. U. S. A. 113, 596–601 (2016).

51. Lewis, J. D. & Izaurflde, E. The Role of the Cap Structure in RNA Processing and Nuclear Export. Eur. J. Biochem. 247, 461–469 (1997).

52. Debarnot, C. et al. Crystallization and diffraction analysis of the SARS coronavirus nsp10-nsp16 complex. Acta Crystallogr. Sect. F Struct. Biol. Cryst. Commun. 67, 404–408 (2011).

53. Viswanathan, T. et al. Structural Basis of RNA Cap Modification by SARS-CoV-2 Coronavirus. bioRxiv Prepr. Serv. Biol. 2020.04.26.061705 (2020) doi:10.1101/2020.04.26.061705.

54. Ananthakrishnan, R., Chard, K., Foster, I. & Tuecke, S. Globus platform-as-a-service for collaborative science applications. Concurr. Comput. Pract. Exp. 27, 290–305 (2015).

55. Chard, K., Tuecke, S. & Foster, I. Efficient and secure transfer, synchronization, and sharing of big data. IEEE Cloud Comput. (2014) doi:10.1109/MCC.2014.52.

56. Clabbers, M. T. B., Gruene, T., Parkhurst, J. M., Abrahams, J. P. & Waterman, D. G. Electron diffraction data processing with DIALS. Acta Crystallogr. Sect. D Struct. Biol. (2018) doi:10.1107/S2059798318007726.

57. Chard, R. et al. FuncX: A Federated Function Serving Fabric for Science. in HPDC 2020 - Proceedings of the 29th International Symposium on High-Performance Parallel and Distributed Computing (2020). doi:10.1145/3369583.3392683.

58. Babuji, Y. et al. Parsl: Pervasive parallel programming in Python. in HPDC 2019-Proceedings of the 28th International Symposium on High-Performance Parallel and Distributed Computing (2019). doi:10.1145/3307681.3325400.

59. Allcock, W. E. et al. Petrel: A Programmatically Accessible Research Data Service. in Proceedings of the Practice and Experience in Advanced Research Computing on Rise of the Machines (learning) (ACM, 2019).

60. Uervirojnangkoorn, M. et al. Enabling X-ray free electron laser crystallography for challenging biological systems from a limited number of crystals. Elife 2015, (2015).

61. Lebedev, A. A., Vagin, A. A. & Murshudov, G. N. Model preparation in MOLREP and examples of model improvement using X-ray data. in Acta Crystallographica Section D: Biological Crystallography vol. 64 33–39 (International Union of Crystallography, 2007).

62. Murshudov, G. N. et al. REFMAC5 for the refinement of macromolecular crystal structures. Acta Crystallogr. Sect. D Biol. Crystallogr. 67, 355–367 (2011).

63. Emsley, P. & Cowtan, K. Coot: Model-building tools for molecular graphics. Acta Crystallogr. Sect. D Biol. Crystallogr. 60, 2126–2132 (2004).

64. Pettersen, E. F. et al. UCSF Chimera - A visualization system for exploratory research and analysis. J. Comput. Chem. 25, 1605–1612 (2004).

65. White, T. A. et al. Crystallographic data processing for free-electron laser sources. Acta Crystallogr. Sect. D Biol. Crystallogr. 69, 1231–1240 (2013).

66. Weiss, M. S. Global indicators of X-ray data quality. J. Appl. Crystallogr. (2001) doi:10.1107/S0021889800018227.

67. Karplus, P. A. & Diederichs, K. Linking crystallographic model and data quality. Science (80-.). 336, 1030–1033 (2012).

68. Davis, I. W., Murray, L. W., Richardson, J. S. & Richardson, D. C. MolProbity: Structure validation and all-atom contact analysis for nucleic acids and their complexes. Nucleic Acids Res. (2004) doi:10.1093/nar/gkh398.

69. Wallace, A. C., Laskowski, R. A. & Thornton, J. M. Ligplot: A program to generate schematic diagrams of protein-ligand interactions. Protein Eng. Des. Sel. (1995) doi:10.1093/protein/8.2.127.

